# Gender-based heterogeneity of FAHFAs in trained runners

**DOI:** 10.1101/2023.06.07.543941

**Authors:** Alisa B. Nelson, Lisa S. Chow, Donald R. Dengel, Meixia Pan, Curtis C. Hughey, Xianlin Han, Patrycja Puchalska, Peter A. Crawford

## Abstract

Fatty acid esters of hydroxy fatty acid (FAHFA) are anti-diabetic and anti-inflammatory lipokines. Recently FAHFAs were also found to predict cardiorespiratory fitness in trained runners. Here we compared the association between circulating FAHFA baseline concentrations and body composition, determined by dual x-ray absorptiometry, in female runners who were lean (BMI < 25 kg/m^2^, n = 6), to those who were overweight (BMI ≥ 25 kg/m^2^, n = 7). We also compared circulating FAHFAs in lean male runners (n = 8) to the same trained lean female (n = 6) runner group. Circulating FAHFAs were increased in females in a manner that was modulated by specific adipose depot sizes, blood glucose, and lean body mass. As expected, circulating FAHFAs were diminished in the overweight group, but, strikingly, in both lean and overweight cohorts, increases in circulating FAHFAs were promoted by increased fat mass, relative to lean mass. These studies suggest multimodal regulation of circulating FAHFAs and raise hypotheses to test endogenous FAHFA dynamic sources and sinks in health and disease, which will be essential for therapeutic target development. Baseline circulating FAHFA concentrations could signal sub-clinical metabolic dysfunction in metabolically healthy obesity.

## Introduction

Fatty acid esters of hydroxy fatty acids (FAHFAs) form a lipid class in which each species is composed of a fatty acyl chain esterified to a hydroxy fatty acid. These lipids serve as signaling lipokines with insulin-sensitizing and anti-inflammatory effects^1-5^. While FAHFA concentrations vary among rodent tissues and tend to be higher in abundance in humans with lower body mass index (BMI) and higher insulin sensitivity, their regulation and mechanisms of action are not yet fully understood^1,6-8^. We previously measured the concentrations of 25 FAHFA species using multidimensional mass spectrometry-based shotgun lipidomics in a cohort of trained runners across a range of BMI to investigate signatures that might identify metabolically healthy obese (MHO) participants^9^. As expected, baseline circulating FAHFAs (participants at rest and after ≥8-hour fast) were negatively associated with BMI and total fat mass. FAHFAs were dynamically regulated during acute aerobic exercise in lean participants, but they were largely unchanged in overweight or obese participants, indicating a concealed effect of obesity in an otherwise MHO group^10^.

We hypothesized that circulating FAHFAs reflect differences in body composition. Therefore, in this study, we investigated relationships between baseline circulating FAHFAs in trained runners relative to dual x-ray absorptiometry (DXA) based body composition measurements.

## Results and Discussion

### Baseline circulating levels of FAHFAs in trained runners are associated with gender, but only in lean individuals

Variability of circulating FAHFAs in the NWT group led us to investigate whether a gender-based (participant self-identified) distinction in baseline circulating FAHFAs could be observed. After pooling all male, female, normal weight trained (NWT, BMI < 25 kg/m^2^), and overweight or obese trained (OWT, BMI ≥ 25 kg/m^2^) participant data per FAHFA species, serum concentrations of many FAHFA species were significantly higher than the mean in female NWT runners compared to male NWT **(Figure 1A)**. Meanwhile, the OWT group did not show any differences in circulating FAHFAs between male and female participants.

**Figure 1.**
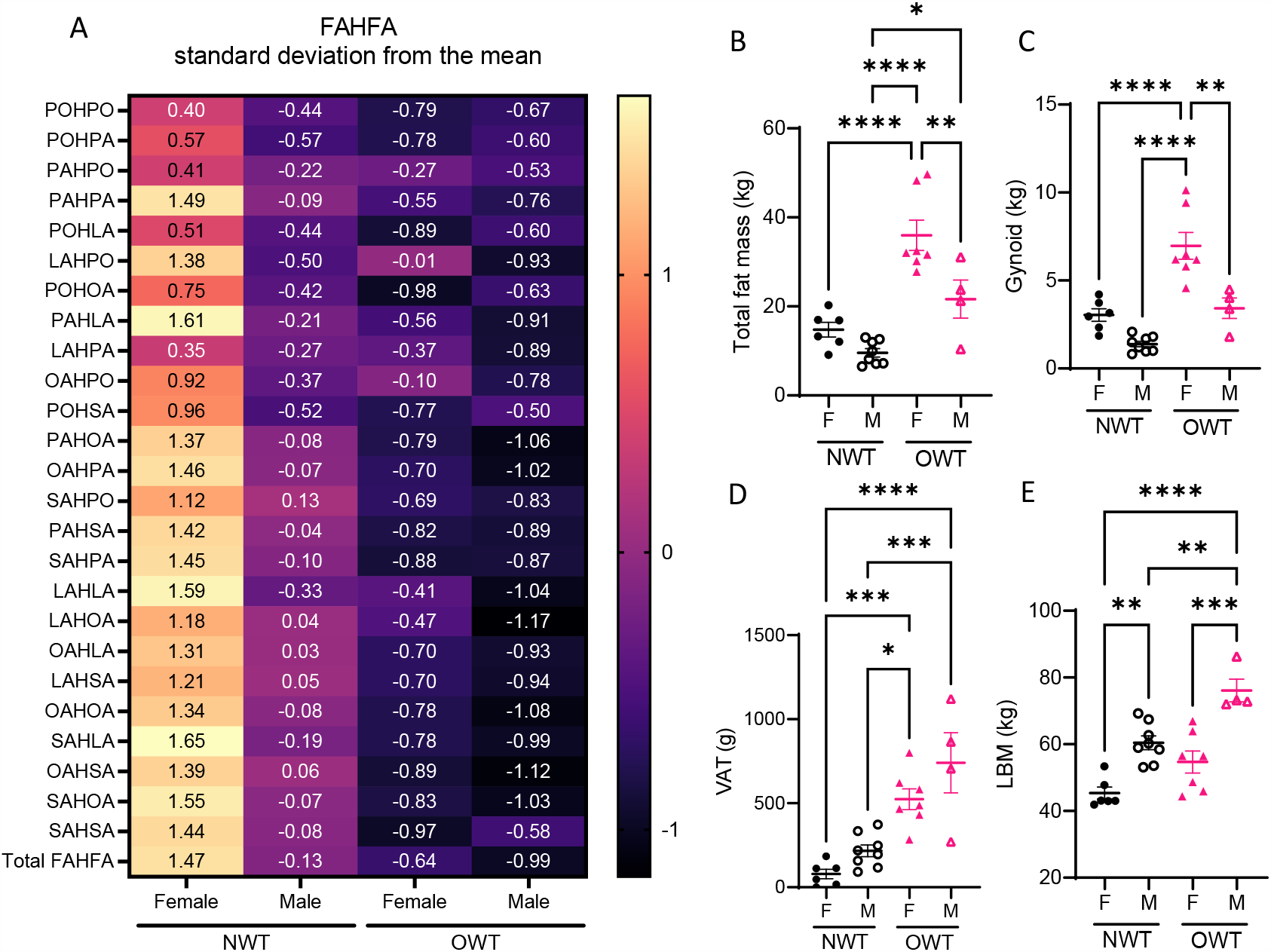
Gender differences in baseline circulating FAHFAs in lean, but not overweight, runners. **(A)** Auto scaled (mean-centered, divided by standard deviation) of circulating concentration of FAHFA species of female and male participants of normal weight-trained (NWT) and overweight and obese trained (OWT) groups. The numerical label indicates deviation from the mean of FAHFA concentration across all participants for each subgroup. **(B)** Total fat mass; **(C)** gynoid fat mass; **(D)** visceral adipose tissue (VAT) fat mass; **(E)** lean body mass (LBM). \**p*<0.05, \*\**p*<0.01, \*\*\**p*<0.001, \*\*\*\**p*<0.0001 using One-way ANOVA with Tukey’s multiple comparisons test. Error bars represent SEM.

Previous work that examined fewer FAHFA species did not identify gender differences in circulating FAHFAs^11^. Additionally, our study involved self-reported trained runners^9^. As we observed both gender and BMI-related differences in circulating FAHFAs, we hypothesized adipose depots may underlie these differences **(Table 1)**. While the OWT group had greater adiposity than the NWT group, OWT females had significantly more total fat mass than OWT males **(Figure 1B)**. OWT females also had greater gynoid fat mass than all other participants **(Figure 1C)**. Visceral adipose tissue (VAT) of the android compartment, is associated with cardiometabolic disease risk^12^. Female participants of the NWT group had the lowest VAT (78.8 ± 28.5g). OWT male participants (740.8 ± 178.3g) had 1.4-fold greater VAT than OWT females (524.0 ± 61.7g), 3.4-fold greater than NWT males (217.1 ± 35.0g), and 9-fold greater compared to NWT females **(Figure 1E)**. Finally, male participants in both NWT and OWT had significantly more lean mass (LBM) than corresponding females **(Figure 1E)**.

**Table 1.** Participant characteristics. SE: standard error; BMI: body mass index; VAT: visceral adipose tissue; SAT: subcutaneous adipose tissue; HOMA-IR: Homeostatic model assessment for insulin resistance; NWT: normal weight trained runners; OWT: overweight or obese trained runners. a: *p-value < 0*.*05 compared to NWT males;* b: *p-value < 0*.*05 compared to NWT females;* c: *p-value < 0*.*05 compared to OWT males*.

|  | NWT (N = 14) |  | OWT (N = 11) |  |
| --- | --- | --- | --- | --- |
| Mean (SE) | Male (N = 8) | Female (N = 6) | Male (N = 4) | Female (N = 7) |
| Age (years) | 30.2 (2.0) | 26.6 (2.5) | 30.3 (3.1) | 33.0 (1.8) |
| BMI (kg/m<sup>2</sup>) | 22.2 (0.6) | 21.5 (0.6) | 29.1 (2.0)<sup>a,b</sup> | 31.5 (2.0)<sup>a,b</sup> |
| Lean body mass (kg) | 60.5 (2.1) | 45.4 (1.8)<sup>a</sup> | 76.1 (3.4)<sup>a,b</sup> | 54.7 (3.3)<sup>c</sup> |
| Total fat mass (kg) | 9.6 (0.9) | 14.8 (4.0) | 21.6 (4.3)<sup>b</sup> | 36.0 (3.4)<sup>a,b,c</sup> |
| Android fat (g) | 569.5 (111.8) | 772.0 (117.9) | 2033 (593.7)<sup>a</sup> | 2853 (371.7)<sup>a,b</sup> |
| VAT (g) | 217.1 (35.0) | 78.8 (28.5) | 740.8 (178.3)<sup>a,b</sup> | 524.0 (61.7)<sup>a,b</sup> |
| SAT (g) | 352.6 (86.9) | 693.2 (114.4) | 1293 (415.9) | 2329 (357.1)<sup>a,b</sup> |
| Gynoid (g) | 1386 (159.7) | 3040 (353.7) | 3425 (586.4) | 6967 (765.7)<sup>a,b,c</sup> |
| HOMA-IR | 0.86 (0.13) | 0.98 (0.25) | 1.97 (0.27)<sup>a</sup> | 1.35 (0.30) |

### FAHFAs differentially associate with body composition

Although FAHFA concentrations have been quantified in numerous rodent tissues, including various adipose depots, liver, and skeletal muscle, little is known about their turnover (*i*.*e*., specific sites for FAHFA production and consumption)^4,8^. Studies investigating enzymes involved in their production suggest FAHFAs originate from adipose tissue with other tissues acting as sites of disposal, but this has yet to be elucidated in physiologically dynamic states^1,3,6,7,13^. Association analyses may lead to hypotheses of FAHFA dynamics, possibly conferring the roles of source and sink to adipose and lean tissues, respectively. Therefore, we employed the Pearson correlation method between fat mass depots and LBM with circulating FAHFAs in (i) female (all NWT + OWT) participants (rationale: females were more dynamic comparing NWT to OWT groups, **Figure 2A**) and (ii) all NWT (male + female) participants (rationale: the gender difference was more pronounced in the NWT group, **Figure 2B**).

**Figure 2.**
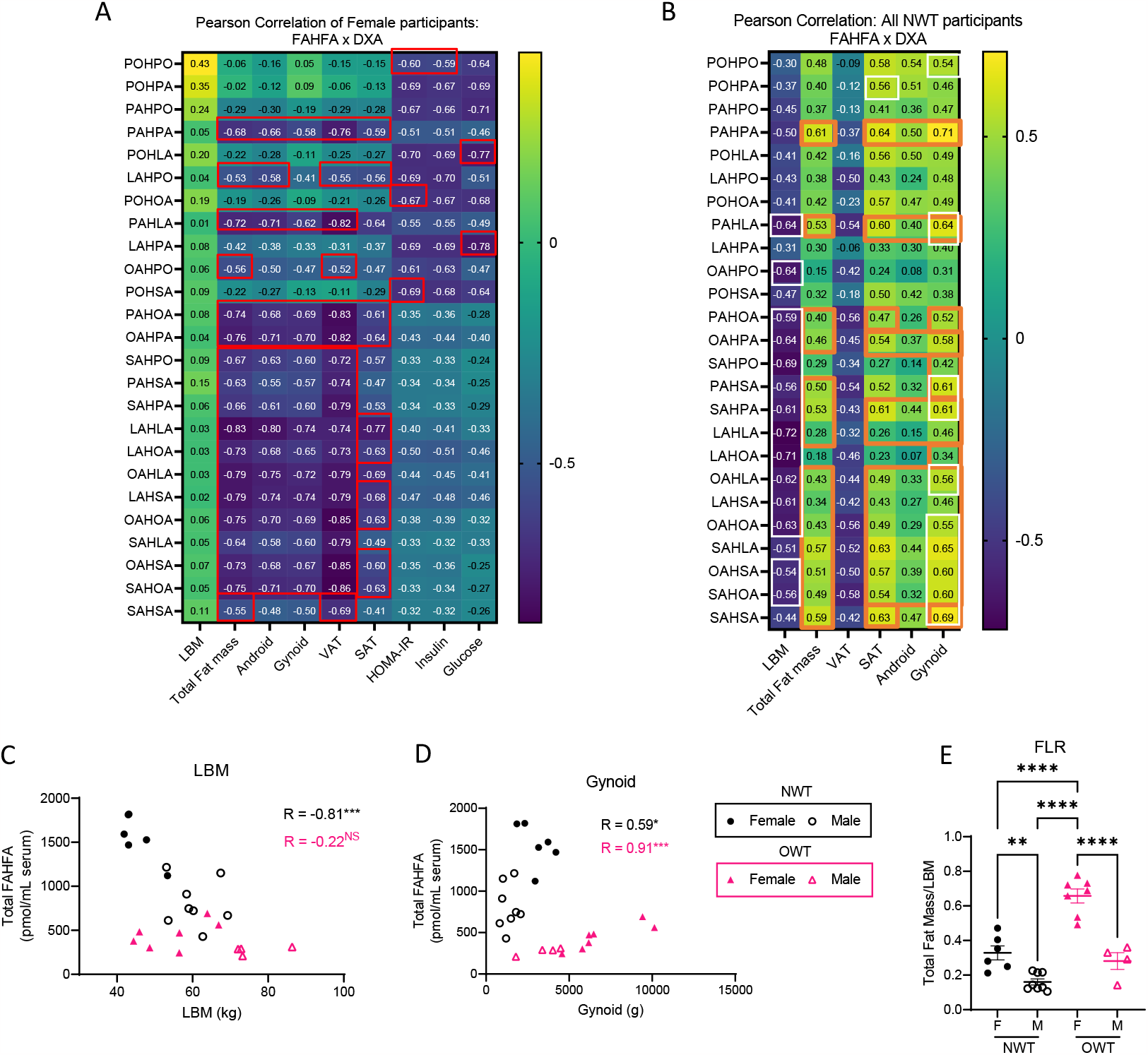
Differential associations between FAHFAs and body composition suggest multimodal influence of FAHFAs by gender and BMI. **(A)** Pearson correlation coefficient of raw circulating concentrations of FAHFA species with DXA measurements, calculated HOMA-IR score, circulating insulin, and circulating glucose for all (NWT + OWT) female participants (n = 13). **(B)** Pearson correlation coefficient of raw circulating concentrations of FAHFA species with DXA measurements for male and female participants (NWT group only, n = 14). |R > 0.5| have a *p* < 0.05; Red box: adj. *p* < 0.05; White box: adj. *p* < 0.1; Orange box: differential correlation between Female and NWT: *p* < 0.01. Scatter plots of **(C)** LBM and **(D)** gynoid fat mass against total circulating FAHFAs; **(E)** total fat to LBM ratio (FLR). NS = not significant, \**p*<0.05, \*\**p*<0.01, \*\*\**p*<0.001 using One-way ANOVA with Tukey’s multiple comparisons test. Error bars represent SEM.

When examining all (NWT + OWT) female participants (n=13) many FAHFA species showed strong negative associations with all fat mass depots (red box indicates those with adj. *p* < 0.05 after correcting for multiple comparisons) **(Figure 2A)**. Significantly strong negative relationships between 18 out of 25 FAHFAs (including the most widely studied FAHFA, palmitic acid ester of hydroxy stearic acid, PAHSA) and total fat mass were observed. No significant differences in the strength of these relationships were seen among specific adipose tissue depots. These findings indicate that in this cohort of exercise-trained females, higher fat mass was associated with lower circulating FAHFAs and does not appear to be associated with a specific depot. Interestingly, five FAHFA species (PAHPO, POHLA, POHOA, LAHPA, and POHSA, but not PAHSA) negatively correlated with circulating insulin, glucose, or HOMA-IR values **(Figure 2A)**. LAHPA showed the strongest negative relationship with fasting glucose (R = -0.78 [95% CI: -0.93 to -0.40], *p* = 0.002). However, none of these five species was negatively associated with fat mass in the lean, trained female participants (LAHPA with Total fat mass: R = -0.38 [95% CI: -0.77 to 0.22], *p* = 0.2)^14,15^. This unexpected relationship suggests these species might be independently regulated by adiposity versus hormone-fuel relationships.

Strikingly, correlation analysis of the NWT (male + female) group (n = 14) shows that 14 out of 25 FAHFA species (including PAHSA) had negative associations with LBM (Pearson correlation coefficient, R < -0.5; white box indicates those with adj. *p* < 0.1 after correcting for multiple comparisons) **(Figure 2B)**. Given prior positive associations of circulating FAHFAs with leanness, this observation, generated in a focused analysis of lean, trained human participants, was unexpected^9,11^. Moreover, relationships to total fat mass, SAT fat (13 of 25 species, R ≥ 0.5), and gynoid fat (13 of 25 species R ≥ 0.5) were positively associated with baseline circulating FAHFA concentrations in the NWT cohort, which was also unexpected. Differential associations (R_Female_ in **Figure 2A** vs R_NWT_ in **Figure 2B**, where orange boxes indicate *p* < 0.01) between circulating FAHFAs and specific fat depot sizes strongly suggest that excess adiposity results in alterations to the regulation of FAHFA turnover (i.e., production and/or consumption) (**Figure 2B**). These differences can be further observed by comparing NWT (n=14) associations to OWT participants (n = 11) **(Figure 2C-F)**. While the NWT group shows a significant inverse relationship between FAHFAs and LBM, the OWT shows no relationship **(Figure 2C)**. Additionally, as in the NWT group, fat mass depots, including the gynoid compartment, show direct relationships with total circulating FAHFAs in the OWT group, but the OWT group carries a highly distinct Pearson correlation coefficient compared to that of NWT participants **(Figure 2D)**.

The distinct clustering of circulating FAHFA distributions by BMI supports the hypothesis that excess adiposity influences regulation of FAHFA turnover. The anthropometric index, total fat to lean mass ratio (FLR), was previously reported to correlate with cardiometabolic disease^16^. In our study population, FLR is highest in OWT females **(Figure 2E)**. Greater FLR in females than males, and higher FAHFA levels in females than BMI-matched males, suggests that increases in circulating FAHFAs are promoted by increased fat mass relative to LBM. This unexpected relationship is evident independently in both lean and overweight groups (**Figure 2B, Figure 2D**). However, when lean and overweight groups are combined, FAHFA concentrations show the expected negative association with body fat (**Figure 2A**). Thus, it is likely that pathogenic adipose expansion, particularly in VAT, negatively influences circulating FAHFA concentrations. These relationships may herald clinical markers of metabolic dysfunction in MHO.

These analyses suggest that circulating FAHFA levels reflect complex regulation influenced by both gender and BMI. Whether these differences are unique to trained humans remains unknown and requires further study to determine how specific compartments dynamically produce and consume FAHFAs in varying states of adiposity and in specific lean and adipose tissue depots. These data also support the notion that select FAHFA lipid species may be independently regulated, and future studies will determine FAHFA lipid species-specific turnover across BMI groups, training status, and gender to parse the multidimensional regulation of circulating FAHFAs. Altogether, these studies raise hypotheses to test endogenous FAHFA dynamic sources and sinks in health and disease, which will be essential for therapeutic target development, and reveal that baseline circulating FAHFA concentrations could signal sub-clinical metabolic dysfunction in MHO.

## Methods and Materials

### Participant details

Overweight trained (OWT; n=11) and normal weight trained (NWT; n=14) participants who self-reported aerobic exercise (3-5 sessions/week) from the Twin Cities metro area were recruited between July 2014 and April 2017. We preferentially recruited participants from recent running events, to ensure that they are capable to complete a prolonged (90 minute) run. Inclusion criteria were: 1) Age 18-40 years, and 2) Regular aerobic exercise, preferably running, at least 3-5 sessions/week. Individuals with 1) Self-reported clinically significant medical issues (for example diabetes, cardiovascular disease, uncontrolled pulmonary disease), 2) abnormal EKG indicating cardiac disease (study EKG performed) and 3) current pregnancy (screening pregnancy test performed) were excluded. Participants were recruited with the goal to achieve similarity in age and sex between the two groups. The University of Minnesota’s Institutional Review Board (IRB) approved the study protocol and methods. All participants provided written informed consent before study participation. Blood was drawn after an 8-hour fast to measure insulin and glucose levels, which were also used to calculate insulin sensitivity, as estimated by homeostatic model assessment for insulin resistance [HOMA-IR: (fasting serum insulin (uU/mL) fasting glucose (mmol/L))/22.5]^14,15^ (14, 15). These samples were also used to measure baseline FAHFA concentrations prior to acute aerobic exercise.

### Dual X-ray Absorptiometry

Lean mass, bone mass, bone mineral content, and fat mass were measured by dual X-ray absorptiometry (DXA) using a GE Healthcare Lunar iDXA (GE Healthcare Lunar, Madison, WI) with Encore software (version 16.2). The DXA scan was performed in the fasted state during a separate visit. Regional measures of trunk, android, gynoid, abdominal subcutaneous adipose tissue (SAT), and visceral adipose tissue (VAT) were made. Trunk fat mass included fat mass from the chest, abdomen, and pelvis region. The android region was defined as the trunk area approximately between the ribs and the pelvis. The upper boundary was set at 20% of the distance between the iliac crest and the base of the skull. The lower boundary was the top of the iliac crest. The gynoid region included the hips and upper thighs, overlapping both the leg and trunk regions. SAT in the android region was determined by examining the X-ray attenuation between the edge of the body and the outer edge of the abdominal cavity^17^. VAT in the android region was calculated by subtracting android SAT from android total fat mass, as previously described^17^.

### Quantitation of FAHFA

Quantitation of FAHFA was previously described^9,18^. Briefly, FAHFA were identified and quantified through multidimensional MS-shotgun lipidomics using appropriate amount of IS 12-PAHSA-d4. Lipids were extracted using a modified Bligh and Dyer protocol and solid phase extraction. Before infusion, FAHFAs were derivatized with N-[4-(Aminomethyl)phenyl]pyridinium (AMPP). Then samples were infused in TSQ triple quadrupole mass spectrometer (Thermo Fisher Scientific) equipped with an automated nanospray device (Triverse Nanomate, Advion Biosciences).

### Statistics

FAHFA concentration was auto scaled. Briefly, the average concentration was computed for each FAHFA species based on all 25 participants. Each sample value was then mean-centered and divided by the standard deviation of each variable. Group differences in body composition, and FLR analyzed using Ordinary one-way ANOVA with Tukey’s multiple comparisons test. Correlation analysis between FAHFA species and body composition were computed using the Pearson correlation in GraphPad Prism v. 9.5.1 for (i) all female participants (n = 13); (ii) all NWT participants (n = 14); (iii) all OWT participants (n = 11) and corrected for multiple comparisons using Benjamini-Hochberg method.

## Acknowledgements

We acknowledge and appreciate those who participated in our study. This work was supported by these funding sources: NIH DK091538, AG069781, DK098203, T32DK007203, P30 AG013319, P30 AG044271.

